# Elemental Analysis of Herbal Food Supplements using ICP-MS for Toxicant and Nutritional Profiling

**DOI:** 10.64898/2026.06.17.733016

**Authors:** Yihunie Hibstie Asres, Manny Mathuthu

## Abstract

Botanical dietary supplements (like wheat, barley, teff, oats, white lupin, pumpkin seed, and chickpeas) may contain trace amounts of toxicants in addition to important micronutrients. Developing and validating a reliable protocol for the simultaneous quantification of Cu, Fe, Zn, Mo, Se, Mn, Pb, Al, Ni, and Cr using a PerkinElmer (NexION^TM^2000 model) quadrupole ICP MS (including a He collision and reaction cell when needed) with closed vessel microwave digestion using (HNO_3_ + H_2_O_2_) was the aim of this study.The method was subsequently utilized in a sample survey, and the outcomes were evaluated against WHO/JECFA standards. From five study regions, twenty-seven farm-collected botanical powder samples representing seven species were acquired. To create one composite per species, field subsamples were cleaned, air dried, ground, and blended (nine subsamples per botanical: three grabs from each of three farms). HNO_3_/H_2_O_2_ was used to digest aliquots (0.250–0.500gm) in closed microwave containers. Internal standards, multi-point external calibration, procedural blanks, verified reference materials, matrix spikes, and duplicates were all used in ICP MS’s multi-element quantitation. Method LODs/LOQs, accuracy (CRM recoveries), and precision (RSD) were calculated.The technique produced low LODs that were suitable for dietary evaluation (typical LOD ranges: Cu, Fe, Zn, Mn, Ni, Cr (0.001–0.01) mg/kg; Mo, Se, Pb, Al (0.002–0.05) mg/kg. For the majority of analytes, within-run RSDs were less than 5%, while CRM recoveries ranged from 88.9 to 110%. The concentrations of essential elements varied greatly (average mg/kg: Fe (280.7±25.6); Zn (6.0±0.541); Cu (2.8±0.269); Mn (398.3±23.8); µgm/kg: Se (0.061±0.006); Mo (1.0 ±0.022). Although some composites approached or exceeded conservative intake thresholds for Pb and Al under high consumption scenarios, toxic elements were generally low (mean mg/kg: Pb (0.062±0.007); Al(185.2±18.5); Ni(1.6±0.163); Cr(1.8±0.171).For the simultaneous nutritional and contaminant profiling of supplements derived from cereals and those not, the validated ICP- MS workflow with microwave HNO_3_ and H_2_O_2_ digestion is suitable. Accurate labeling and consumer safety can be supported by routine screening and supply chain controls.

## Introduction

Herbal food supplements, particularly those derived from cereals and non-cereals (wheat, barley, teff, oats, white lupin, pumpkin seed, and chickpea), are increasingly used to improve micronutrient intake and overall dietary quality[1,2]. Essential trace metals like Cu, Fe, Zn, Mo, Se, and Mn are found naturally in these plant materials and support vital physiological processes [2-4]. In parallel, herbal supplements may also contain trace toxicants, notably Pb, Al, Ni and Cr, with their concentrations influenced by factors including raw-material provenance, agricultural practices, and post-harvest processing or handling conditions[1]. Consequently, an analytical strategy capable of simultaneously quantifying both nutritional elements and potential contaminants is critical for evidence-based safety screening and nutritional profiling [3].

A major public-health and regulatory challenge is the limited availability of multi-element datasets that are sufficiently sensitive, quality-assured, and comparable across essential and contaminant species for botanical food supplements [5,6]. Many published studies focus either on essential nutrients only or on a narrow set of contaminants, which restricts the usefulness of these data for exposure-relevant risk screening and integrated nutritional assessment [7]. Additionally, the natural heterogeneity of botanical matrices, varying by botanical species, sampling region, and processing lots, can lead to substantial differences in elemental composition [6]. This variability, combined with inconsistencies in analytical workflows across studies, complicates interpretation and limits cross-comparability [8].

Standardized and validated analytical techniques that can quantify a wide range of elements at low concentrations and produce results that can be interpreted using established guidance frameworks, like WHO/JECFA-related intake metrics, are therefore practically needed [9, 10]. A key research gap remains the lack of validated, multi-element analytical data for herbal food supplements that cover both essential nutrients and trace toxicants using robust, quality-controlled workflows[11]. Many studies do not report comprehensive method validation parameters (limits of detection/quantification, recovery, and precision), which reduces confidence in the results and hinders reliable comparison across product types. Furthermore, defensible exposure-relevant data are particularly important for heterogeneous botanical ingredients [11].

To bridge this gap, the current study set out to develop and evaluate a dependable ICP-MS methodology for the simultaneous elemental profiling of herbal dietary supplements manufactured from cereal and non-cereal botanicals [12,13].A quadrupole ICP-MS method (NexION*^TM^*2000 model) was developed and validated and coupled with closed-vessel microwave digestion using (HNO_3_ + H_2_O_2_) [14].The method was used to quantify Cu, Fe, Zn, Mo, Se, Mn, Pb, Al, Ni, and Cr in farm-collected botanical seed flour composites [4,11]. Following method validation, the workflow was applied to generate multi-element data for a survey of samples and to support interpretation of results in relation to WHO/JECFA guidance and relevant dietary contexts [10,15].

The novelty and strength of this study lie in: (i) integrating a broad panel of nutritional micronutrients and targeted trace toxicants within a single analytical framework; (ii) using validated microwave digestion and ICP-MS quantification to achieve low detection limits suitable for dietary assessment; and (iii) strengthening methodological credibility through determination of LOD/LOQ, certified reference material (CRM) recoveries, and precision (RSD). Collectively, this work provides a validated analytical platform that can enhance safety screening and support accurate nutritional labelling of herbal food supplements derived from cereal and legume botanicals [12].

## Materials and Methods

### Study design

A study design is an experimental analytical cross-sectional approach in which herbal food supplement samples are collected from different locations, coded, and stored to maintain sample integrity.ICP-MS is used to detect the amounts of target metals in these samples following microwave digestion [16,17]. The elemental concentration data are then analyzed to produce nutritional profiling and toxicant-related (carcinogenic and non-carcinogenic health) insights using quantitative modeling of metal levels across the sampled products [16]. The obtained elemental concentration data are then analyzed using quantitative modeling of metal levels across the selected goods to produce nutritional profiling and toxicant-related (carcinogenic and non-carcinogenic health) insights [16].

### Study area

East Gojjam, a portion of the broader mountainous region known for the Choke mountains and the Blue Nile Gorges, is where the study was conducted.This region has a variety of topographical features, including plateaus, escarpments, plains, gorges, and deeply carved valleys. Elevations in the area range from approximately 2800 to 4100 meters above sea level [18]. The region is also dominated by dry and semi-arid climatic conditions in which cereal and non-cereal production are largely rain-fed, and climate-related changes are expected to affect agricultural systems through processes such as acidification [1].Each sampling location was assigned a numeric code to ensure systematic sample collection and traceability. In order to protect sample integrity throughout the analytical process, samples were handled consistently, labeled using coding before laboratory analysis, and maintained in accordance with the specified preservation technique[19]. National geography mapmarker software was used to create maps of East Gojjam.

### Sample collection and preparation

Purposeful sampling was used to select representative samples of *Cicer arietinum* (chickpea), *Lupinus albus* (lupin), *Eragrostis teff* (teff), *Avena sativa* (oats), *Triticum aestivum* (wheat), *Cucurbita* (pumpkin), and *Hordeum vulgare* (barley) based on expert experience and prior knowledge (**Fig. 1**) [17, 20, 21]. Clean, empty plastic bags were used for collection in order to reduce contamination. Before being ground into a uniform powder and combined using an agate mortar and pestle, the collected samples were air-dried as needed [18,21].

**Fig. 1.**
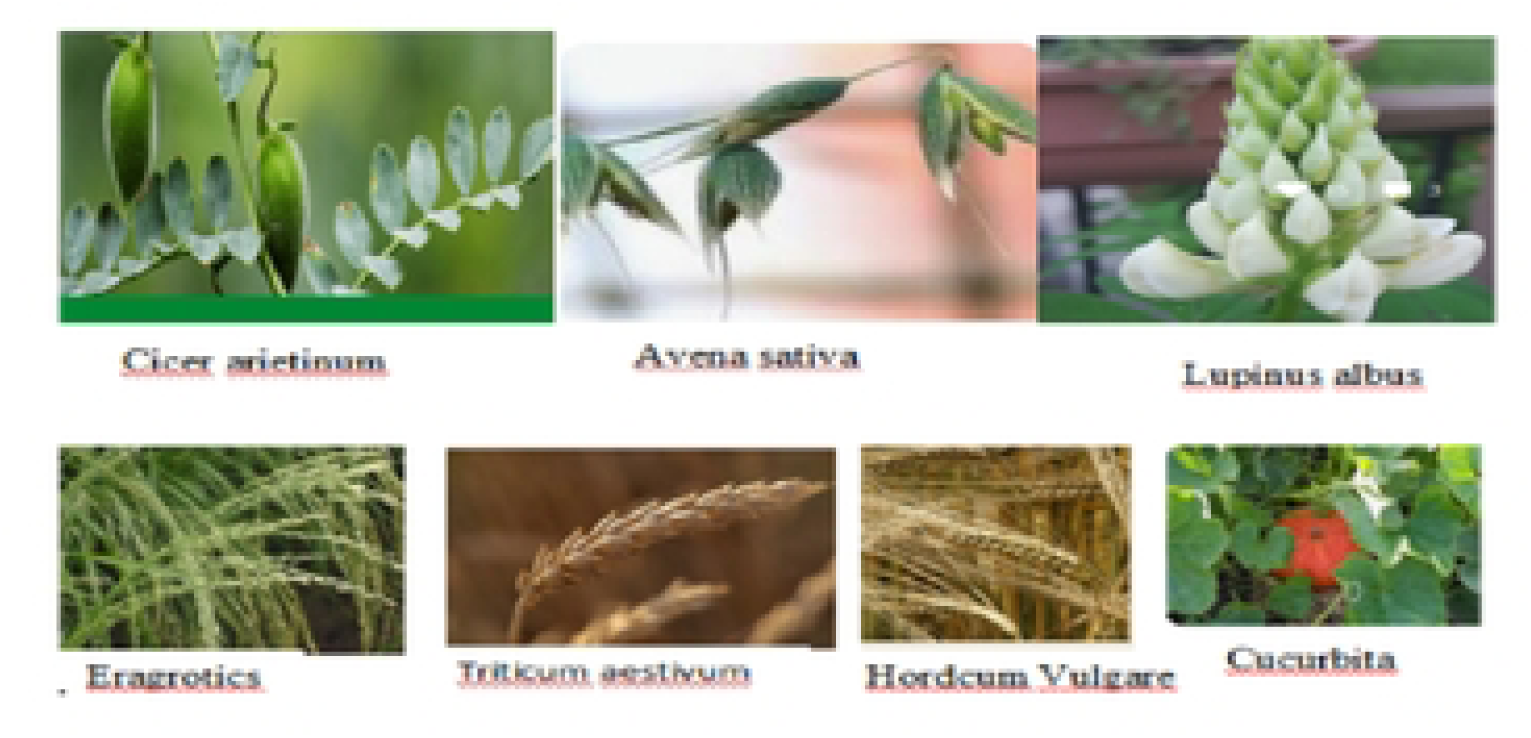
Available cereals and non-cereals in their agricultural area. Source: Own elaboration

To reduce exogenous contamination prior to chemical analysis, the raw materials were rinsed with tap water, then processed through a 100 µm filter to obtain uniform particle size [22]. The homogenized powders were transferred into appropriately labeled plastic containers and stored prior to ICP-MS analysis[23].

### Legality of sample transportation and compliance

All plant materials were handled and documented in accordance with national and institutional requirements to ensure ethical compliance, traceability, and legal transfer. Powdered samples were prepared using voucher specimens of ethnobotanical medicinal plants referenced from the Flora of Ethiopia and Eritrea [24,25].

The specimens were sent to the Center for Applied Radiation Science and Technology (CARST) in South Africa for study after being properly packed, labeled, and sealed by the Ethiopian Biodiversity Institute in Addis Ababa. The receiving laboratory ensures appropriate chain-of-custody and integrity of the submitted samples by processing biological materials in accordance with institutional standard operating procedures and ICP-MS analysis [17,23].

### Microwave Digestion Parameters + Procedure (ICP-MS Sample Preparation)

Before being placed in a tetrafluoroethylene (PTFE) microwave digestion vessel, each herbal food supplement sample (0.30 gm) or CRM (0.30 gm) was carefully weighed. After that, 8 mL of nitric acid (HNO₃, 70%) and 2 mL of hydrogen peroxide (H2O₂, 30%) were added to the vessel.The contents were gently swirled and left for approximately ten minutes to promote the organic matrix. The most crucial elements in the microwave digestion process were determined to be the factor (HNO3 + H2O2) with (70%, 30%), respectively, and the digestion duration [14].To guarantee that the sample was completely transferred to the bottom of the vessel, the vessel was washed with acid. To encourage the organic matrix’s first digestion, the contents were gently swirled and left to stand for around ten minutes before being sealed[14].

The Titan MPS*^TM^* microwave pressure digestion system was used to carry out microwave digestion under the following program (as target settings): three temperature phases with a constant pressure limit of 30 bar. There were two stages: one for 190°C (ramp time 3 minutes, hold time 20 minutes, power 100%) and one for 160°C (ramp time 5 minutes, hold time 5 minutes, power 90%). To finish the digestion and cool the vessels to a safer temperature, the last step was set to 50°C (ramp time 1 min, hold time 1 min, power 0%). The vessels were allowed to cool until they were warm to the touch before being opened in order to minimize foaming and possible sample loss. After the digests were quantitatively transferred into 25 mL polyethylene volumetric flasks and diluted to volume with ultrapure water, ICP-MS analysis was carried out on all produced solutions.

Reagent blanks were created by processing identical volumes of acids under the same conditions without sample material[16]. Ultrapure water was used to dilute the digests to volume after they were quantitatively placed into 25 mL polyethylene volumetric flasks. The same amounts of acids were treated under the same conditions without any sample material to produce reagent blanks[16].

To quantify metals, the digested samples (as shown in **Fig.2**) were diluted in one to ten (1:10), and then in one to hundred (1:100) with distilled water and fed onto an autosampler of an inductively coupled plasma mass spectrometer (NexNOX*^TM^* 2000 model)[26, 23].

**Fig. 2.**
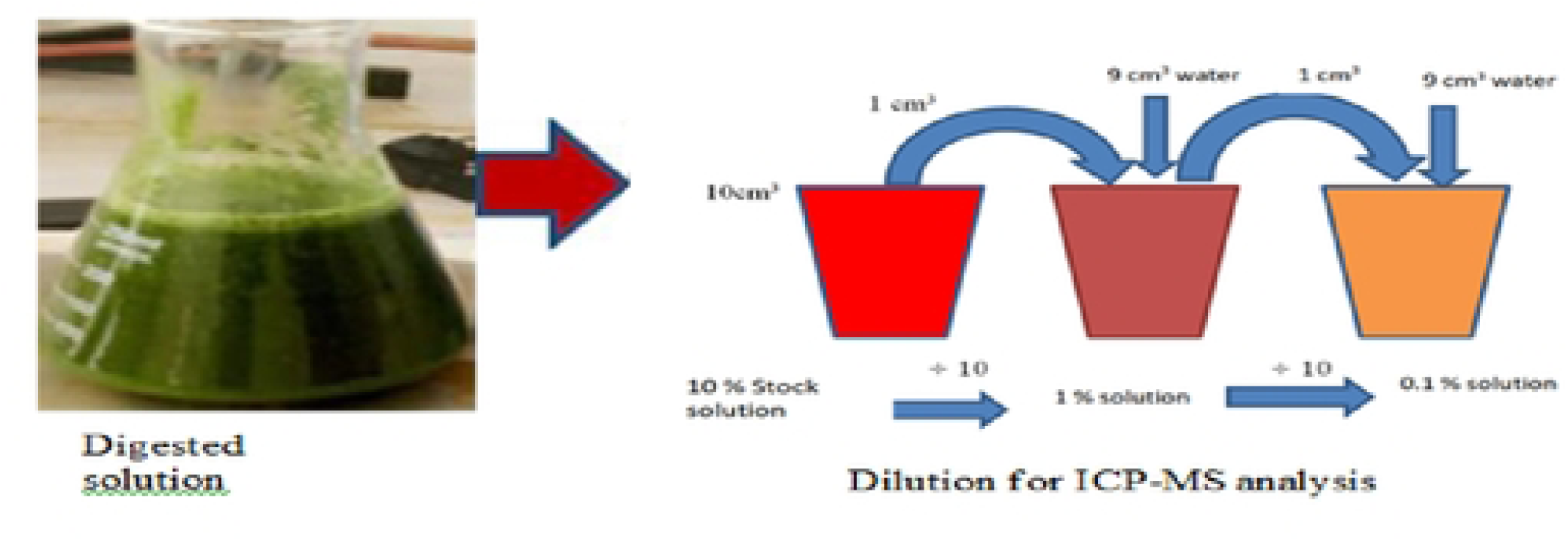
Sample Dilution for Analytical Element Findings. *S*ource: Personal explanation

ICP-MS was used to determine Fe, Zn, Cu, Mn, Se, Mo, Pb, Al, Ni, and Cr. Argon (99.9%) was used as an auxiliary gas for nebulization and plasma generation[26, 27].

According to (**Fig.3)**, the liquid sample is nebulized by argon into fine aerosol droplets and transported through the spray chamber into the ICP torch. In the plasma, droplets are rapidly desolvated (dried), vaporized, and ionized [26, 27]. To increase transport efficiency, extra droplets are removed from the spray chamber. The resulting ions are then moved to the MS interface for examination.

**Fig. 3.**
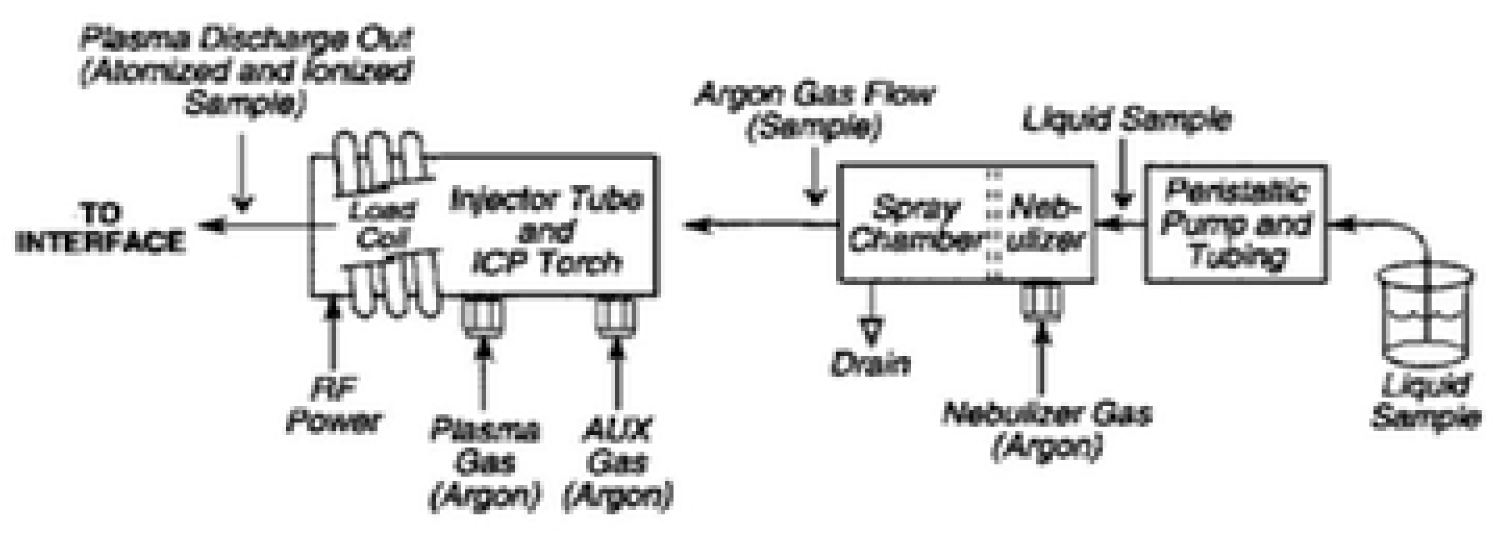
Schematic Representation of an ICP Source Inside an ICP-MS. Source: Personal explanation

Blank samples were analyzed and deducted from the sample readings prior to the computation of the results [26, 27]. The limits of detection (LOD) were computed three times the blank sample standard deviation. The limits of quantitation (LOQ) were computed using the average sample volume and the total volume analyzed. Each sample was measured three times before the average value was utilized. In element analysis, CRMs were employed for both precision and accuracy [28, 29].The blank sample standard deviation was multiplied by three to find the limits of detection (LOD). The average sample volume and the total volume analyzed were used to compute the limits of quantification (LOQ). Before using the average value, each sample was measured three times. CRMs were used to assess element analysis accuracy and precision [28, 29].

### Data Validation

The data was validated using a linearity analysis of calibration curves and recoveries (standard and spike recovery). Each metal’s intensity vs. concentration calibration curves were calculated based on its working standards. The metals’ coefficients of determination (R^2^) ranged from 0.9993 to 0.9999, and their limits of quantification (LOQs) ranged from 0.01 ppm (Cr) to 0.265 ppm (Fe).

Using the same sample calibration, the constructed standards were run once again; the minimum standard recovery for selenium, chromium, and molybdinum was 70%. Calibration curves for intensity/response vs. concentration were created for each metal based on its working standards. The metals under investigation had coefficients of determination (R^2^) ranging from (0.9993-0.9999).

The metals’ limits of quantification (LOQs) ranged from 0.01 ppm (Cr) to 0.265 ppm (Fe)[19]. The generated standards were run again using the same sample calibration. The lowest standard recovery for selenium was 70%, whereas the highest recovery for chromium, molybdinum, and selenium was 110%. To monitor the spike recovery, a known quantity of the representative addition was made for each metal’s standard. Lead had the greatest recovery of 115%, whereas Manganes had the lowest spike recovery of 88.9%.

### Health Risk Assessment

Evaluating the risk to human health includes calculating the relevant indicators (HQ, HI, and ILCR) [19,30,31].

### Non-carcinogenic risk to health

Human health may be negatively impacted by prolonged exposure to a single metal. To forecast detrimental impacts on human health, risk assessments were employed[19,30]. Two types of risk assessment models are non-carcinogenic health risk evaluations. To evaluate non-carcinogenic health risk, the HQ and HI values were computed [32]. The risk level computation is crucial to the risk assessment models [19, 30]. After determining the ingestion rate by the metal content of the sample, the estimated daily intake (EDI) is calculated by dividing the result by body weight [33]. To predict negative effects on human health, risk assessments were employed.Non-carcinogenic health risk was evaluated by computing the HQ and HI values [32]. The risk assessment models require the estimation of the risk level. The estimated daily intake (EDI), which is determined by multiplying the ingestion rate by the metal content in the sample and dividing the result by body weight, indicates the daily consumption of dangerous metals [33].

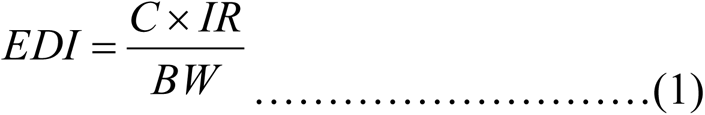

where C (ppm) is the quantity of hazardous metals in herbal dietary supplements, IR (kg/day) is the ingestion rate (20 gm/person/day) or 0.02 kg/person/day to compute, and BW (kg) is the body weight (70 kg).

Chronic daily intake (CDI) is the term used to describe long-term exposure to toxic metals [32, 33]. The current study only considers oral intake.

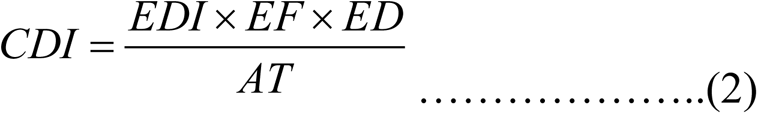

where CDI (ppm/day) is the chronic daily intake, EF (days/year) is the exposure frequency, ED (years) is the exposure duration, and AT (EDx365) is the average time. The CDI to RfD ratio is the HQ for the metal of interest [30, 33]. The CDI to RfD ratio for the metal of interest is represented by the HQ [33].

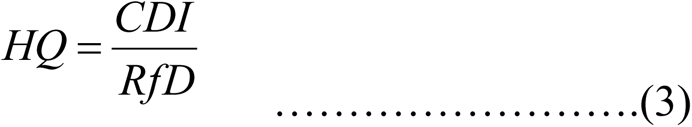

where RfD is the oral reference dosage that permits a person to sustain the exposure level for a long period of time without experiencing any adverse side effects.

Mn (0.14 ppm/day), Fe (0.8 ppm/day), Cu (0.04 pp/day), Ni (0.02 ppm/day), Pb (0.004 ppm/day), Cr (0.003 ppm/day), Zn (0.3 ppm/day), Se (0.005 ppm/day), Mo (0.005 ppm/day), and Al (1.0 ppm/day) [19,30]. If HQ is less than 1 (the risk is deemed low), there is no reason for non-carcinogenic concern [32]. When numerous hazardous metals are exposed, HI is expected to have an extra effect [19, 30]. The total of each metal’s hazard index values is used to calculate the danger index [32, 33].

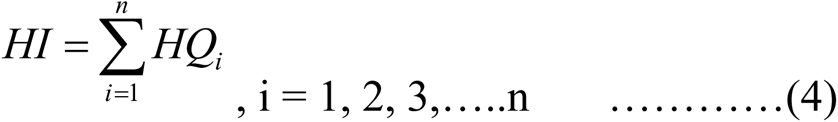

The potential for negative impacts on human health is indicated by an HI value larger than 1.

### Carcinogenic Health Risk

Incremental lifetime cancer risk, or ILCR, is the chance that a person would eventually develop any type of cancer as a result of years of daily exposure to a particular daily dosage of a carcinogenic agent [34].

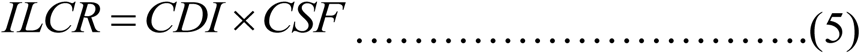

where CSF is the lifetime average risk of one mg/kg/day of cancer-causing metal.

The literature was used to calculate the CSF values for Ni (0.84 ppm/day), Pb (0.0085 ppm/day), and Cr (0.5 ppm/day) [19,30]. The allowable ILCR threshold for a single carcinogenic element is 10^-6^.The allowed ILCR threshold for a single carcinogenic element is 10^-6^. Ni and Cr in herbal food supplements are often higher than the allowable limit (1x10^-6^ to 1x10^-4^)[19,30].

### Statistical analysis

Microsoft Office Excel 2010 was used to tabulate and compile research data. The same program was used for additional data processing, calculations, and storage[19]. In the instance of samples, a histogram created using Origin professional statistical software showed the results of three independent replications (n = 3).The corresponding uncertainty was displayed in the data using positive and negative standard deviations.

## Results and Discussion

### Evaluation of heavy metal health risks

One of the most important techniques is to assess health risks using the estimated daily intake (EDI) of heavy metal pollution [32]. It considers the body weight of those exposed as well as the frequency and duration of exposure [16]. Generally speaking, the average daily dietary consumption determines the health risk associated with metal pollution. The EDI for Pb, Cr, Al, and Ni met the samples’ allowable daily intake reference values [32]. For a sample of wheat, pumpkin, and *Lupines albus*, the EDI for Cr was discovered to be greater than the suggested daily consumption reference limit [12].

HQ was used to determine the non-carcinogenic risk of heavy metals in the herbal dietary supplements.The results are displayed in (**Table 1)** [32]. For non-carcinogenic effects, HQ measures the long-term exposure to heavy metal pollution seen in herbal dietary supplements [14]. HQ values varied from (0.0082-0.5254) ppm/day for Mn, (0.0033-0.2514) ppm/day for Mo, less than (0.0033-0.378) ppm/day for Se, (0.0189-0.1071) ppm/day for Al, (0.0245-0.181) ppm/day for Fe, (0.0081-0.0657) ppm/day for Zn, (0.0011-0.0329) ppm/day for Cu, (0.0743-0.2762) ppm/day for Cr, and (0.003-009) ppm/day for Pb. Heavy metals like Mn, Mo, Se, Al, Fe, Ni, Zn, Cu, Cr, and Pb did not pose any non-carcinogenic health hazards in any of the herbal dietary supplement samples if (HQ < 1)[32]. Customers who are exposed are deemed safe if the HQ value is less than 1. A level of concern or a health risk is indicated if the HQ value is equal to or higher than 1 [19, 30]. According to the results, there are no health hazards associated with long-term consumption of these herbal food supplements because the HQ values for Pb, Al, Ni, and Cr were all less than 1 [31, 32].

**Table 1:** Metals’ hazard index (HI) and hazard quotient (HQ) when ingesting herbal dietary supplements.

| Sample code | HQ |  |  |  |  |  |  |  |  |  | HI |
| --- | --- | --- | --- | --- | --- | --- | --- | --- | --- | --- | --- |
|  | Fe | Zn | Se | Mn | Cu | Mo | Al | Pb | Ni | Cr |  |
| Wheat | 0.181 | 0.011 | 0.378 | 0.048 | 0.0329 | 0.0269 | 0.0774 | 0.0035 | 0.0414 | 0.2476 | 0.7073 |
| Chickpea | 0.0245 | 0.0012 | 0.0015 | 0.0082 | 0.0043 | 0.0075 | 0.0207 | 0.0027 | 0.0092 | 0.1333 | 0.2128 |
| Oats | 0.1444 | 0.0044 | 0.005 | 0.0294 | 0.01 | 0.0118 | 0.039 | 0.0041 | 0.0081 | 0.1429 | 0.999 |
| Teff Red | 0.1042 | 0.0068 | 0.0046 | 0.0888 | 0.0221 | 0.0206 | 0.0745 | 0.0081 | 0.0087 | 0.1333 | 0.4717 |
| Teff White | 0.0514 | 0.0034 | 0.0025 | 0.0371 | 0.0136 | 0.0074 | 0.0189 | 0.0003 | 0.01 | 0.1238 | 0.2684 |
| Black Bareley | 0.0576 | 0.0032 | 0.0032 | 0.0767 | 0.0157 | 0.0037 | 0.0354 | 0.009 | 0.0214 | 0.1524 | 0.3783 |
| WhiteBareley | 0.0329 | 0.0034 | 0.0019 | 0.0245 | 0.0114 | 0.0087 | 0.0232 | 0.0029 | 0.0087 | 0.0743 | 0.1919 |
| LupinusAlbus | 0.1115 | 0.0077 | 0.0054 | 0.5254 | 0.03 | 0.2514 | 0.0798 | 0.0039 | 0.0657 | 0.2762 | 1.4 |
| Pumpkin | 0.114 | 0.0057 | 0.0033 | 0.308 | 0.025 | 0.0033 | 0.1071 | 0.0056 | 0.0357 | 0.2286 | 0.8363 |

The cumulative effect of heavy metal pollution in a particular herbal dietary supplement does not pose a lifetime risk to an adult’s health if the samples’ HI values were less than 1 [32]. However, the exposed population may be at risk for health problems if the HQ is equal to or more than 1 [19, 30].

The cumulative impacts of the heavy metal pollutants in a specific herbal dietary supplement do not provide a long-term health danger to adults if the HI values for samples were less than 1 [14].

Since Cr is toxic and deadly to humans even at lower concentrations, the HI for sample Lupinus albus (HI = 1.4) was more than 1, most likely as a result of a high daily consumption of Cr in this herbal dietary supplement [19].This implies that prolonged exposure to this sample may have detrimental health effects that are not cancer-causing, and quick action is needed to lower the metal levels in the formulation to shield the customer from a possible health danger[32]. This suggests that prolonged exposure to this sample may have negative health impacts that are not cancer-causing, and prompt action is required to lower the metal levels in the formulation to shield the customer from a potential health danger. Even in smaller quantities, Cr is poisonous and deadly to humans [19, 30].

### Risk to Carcinogenic Health

The carcinogenic risks for Pb, Cr, and Ni (**Table2**) were also developed because the metals in issue may contribute to anxiety that is both carcinogenic and non-carcinogenic [32]. The carcinogenic risk was evaluated using the ILCR; values of Pb, Cr, and Ni ranged from 9.7x10^-9^ to 1.2x10^-7^, 1.9x10^-4^ to 4.1x10^-4^, and 1.4x10^-4^ to 1.1x10^-3^, respectively (**Table 2**). Nine of the samples evaluated for Pb, Cr, and Ni had carcinogenic anxiety (ILCR > 10^-6^)[ 19,30].

**Table 2:** Metals’ incremental lifetime cancer risk (ILCR) when ingesting herbal food supplements.

| Sample Code | ILCR |  |  |
| --- | --- | --- | --- |
|  | Pb | Ni | Cr |
| Wheat | $1.2 \times 10^{-7}$ | $7.0 \times 10^{-4}$ | $3.7 \times 10^{-4}$ |
| Chickpea | $9.2 \times 10^{-8}$ | $1.5 \times 10^{-4}$ | $2.0 \times 10^{-4}$ |
| Oats | $1.4 \times 10^{-7}$ | $1.4 \times 10^{-4}$ | $2.1 \times 10^{-4}$ |
| TeffRed | $2.8 \times 10^{-7}$ | $1.5 \times 10^{-4}$ | $2.0 \times 10^{-4}$ |
| Teff White | $9.7 \times 10^{-9}$ | $1.7 \times 10^{-4}$ | $1.9 \times 10^{-4}$ |
| BlackBareley | $3.1 \times 10^{-7}$ | $3.6 \times 10^{-4}$ | $2.3 \times 10^{-4}$ |
| WhiteBareley | $9.5 \times 10^{-8}$ | $1.5 \times 10^{-4}$ | $1.9 \times 10^{-4}$ |
| LupinusAlbus | $1.3 \times 10^{-7}$ | $1.1 \times 10^{-3}$ | $4.1 \times 10^{-4}$ |
| Pumpkin | $1.9 \times 10^{-7}$ | $6 \times 10^{-4}$ | $3.4 \times 10^{-4}$ |

### Heavy metal analysis of herbal dietary supplements

The current study’s findings showed that the nine herbal dietary supplements that are often utilized in East Gojjam included varying concentrations of heavy metals (Pb, Al, Ni, and Cr) [12]. **Table 3** displays the typical level of heavy metals in common herbal dietary supplements [12]. Lead levels in samples of herbal dietary supplements ranged from 0.004±0.004 mg/kg to 0.126±0.013 mg/kg (**Table 3**). The samples utilized by traditional healers to treat sexual impotence and hypertension had the highest and lowest quantities, respectively [35]. One of the most dangerous metals, lead, can cause reproductive abnormalities, gastrointestinal disorders, brain and kidney damage, hearing and visual problems, and poor muscle coordination over time [35, 36].

**Table 3:** Pb (mg/kg), Al (mg/kg), Ni (mg/kg), and Cr (mg/kg) mean concentrations (mean ± SD) in East Gojjam herbal food supplements (n = 3)

| Food Samples | Harmful trace elements |  |  |  |
| --- | --- | --- | --- | --- |
|  | Pb(mg/kg) | Al(mg/kg) | Ni(mg/kg) | Cr(mg/kg) |
| Wheat | 0.049±0.005 | 271.8±27.2 | 2.9±0.29 | 2.6±0.26 |
| Chickpea | 0.038±0.004 | 72.2±7.2 | 0.642±0.06 | 1.4±0.14 |
| Oats | 0.057±0.006 | 136.6±13.7 | 0.57±0.057 | 1.5±0.15 |
| RedTeff | 0.114±0.01 | 260.6±26.1 | 0.610±0.06 | 1.4±0.14 |
| WhiteTeff | 0.004±0.004 | 66.1±6.6 | 0.702±0.07 | 1.3±0.13 |
| BlackBarley | 0.126±0.013 | 123.9±12.4 | 1.5±0.15 | 1.6±0.16 |
| WhiteBarley | 0.039±0.004 | 81.3±8.1 | 0.608±0.06 | 1.3±0.13 |
| LupinusAlbus | 0.054±0.005 | 279.2±27.9 | 4.6±0.46 | 2.9±0.19 |
| Pumpkin | 0.079±0.008 | 374.9±37.5 | 2.5±0.25 | 2.4±0.24 |

The highest level of lead permitted by the WHO in conventional herbal dietary supplements. All of the analyzed samples were within this permissible range [30]. One of the most hazardous metals is lead; prolonged exposure can cause abnormalities in reproduction, gastrointestinal problems, hearing and visual problems, brain and kidney damage, and poor muscle coordination [35, 36]. Patients who consume medicinal herbs containing even small levels of lead over an extended period of time may be at risk for chronic lead toxicity due to lead bioaccumulation in biological tissues, and they should be closely watched for any indications of lead placement [35].

The concentration of chromium in this study ranged from 1.3±0.13 mg/kg to 2.6±0.26 mg/kg (**Table 3**). The wheat sample had the highest content, while the white barley and white teff samples had the lowest[37]. The herbal dietary supplement’s chromium concentration exceeded the WHO-recommended permissible limits of 2 mg/kg [30].This may be due to the fact that, even at lower doses, the raw ingredients utilized to collect the supplements are poisonous and lethal[30]. The white barley and white teff samples had the least amount, whereas the wheat sample had the most[12]. The amount of chromium in the herbal dietary supplement exceeded the 2 mg/kg WHO-recommended permitted limits [30].This may result from the production of herbal dietary supplements using contaminated raw materials [14].

The average daily dietary intake of chromium, which is poisonous and deadly to humans even at lower amounts, determines the overall health risk of metal poisoning [32, 38].

The result of the study indicated that the nine commonly consumed herbal food supplements in East Gojjam contained heavy metals (Pb, Al, Ni and Cr) in varying concentrations[14]. The mean concentrations (mean ± SD) of heavy metals, lead (Pb), aluminum (Al), nickel (Ni) and chromium (Cr) in herbal food supplements in East Gojjam are presented in (**Table3)**. The results show a clear variation between different herbal food supplements[38].Among the hazardous metals examined, aluminum was the most prevalent, with the highest concentrations found in samples of pumpkin (374.9 ± 37.5) and Lupinus albus (279.2 ± 27.9) mg/kg, and the lowest concentrations found in white teff (66.1 ± 6.6) mg/kg. All of the samples had comparatively low Pb concentrations, ranging from (0.004 ± 0.004) mg/kg for white teff to (0.126 ± 0.013) mg/kg for black barley. Lupinus albus had the greatest value of nickel (Ni) (4.6 ± 0.46 mg/kg), whereas oats had the lowest value (0.57 ± 0.057 mg/kg).White teff and white barley had amounts of (1.3 ± 0.13) mg/kg of chromium (Cr), while Lupinus albus had concentrations of (2.9 ± 0.19) mg/kg.

Overall, the toxicant profile obtained from these ICP-MS measurements indicates that the studied herbal food supplements differ substantially in their potential exposure to hazardous elements[38]. Such information is important for safety assessment and for interpreting the nutritional benefits of East Gojjam herbal food supplements alongside possible heavy-metal-related health risks[38]. In particular, samples with higher Al, Ni, and Cr values (pumpkin and *Lupinus albus*) may require additional attention for food safety screening and risk management[39].

**Table 4** summarizes the maximum, minimum, and overall average concentrations of Pb, Al, Ni, and Cr across the investigated herbal food supplements, highlighting the variability among food types and providing a basis for nutritional and safety evaluation.

**Table 4:**
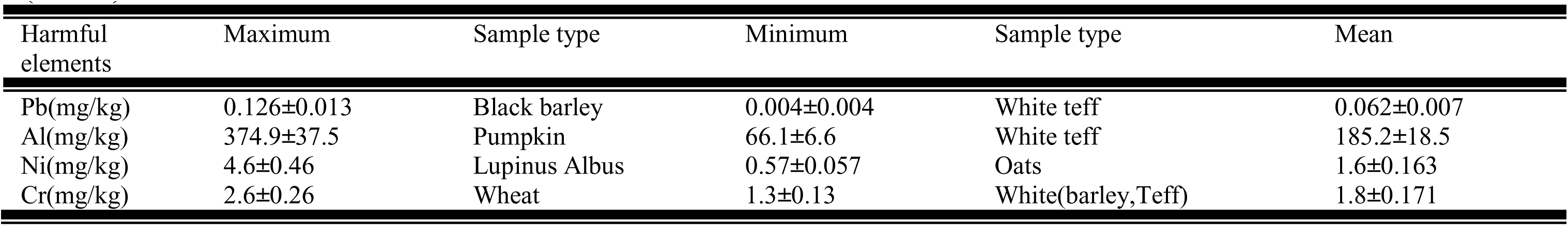
East Gojjam herbal food supplements’ maximum, minimum, and average concentrations (mean ± SD, mg/kg) of Pb (mg/kg), Al (mg/kg), Ni (mg/kg), and Cr (mg/kg) (n = 3)

| Harmful elements | Maximum | Sample type | Minimum | Sample type | Mean |
| --- | --- | --- | --- | --- | --- |
| Pb(mg/kg) | 0.126±0.013 | Black barley | 0.004±0.004 | White teff | 0.062±0.007 |
| Al(mg/kg) | 374.9±37.5 | Pumpkin | 66.1±6.6 | White teff | 185.2±18.5 |
| Ni(mg/kg) | 4.6±0.46 | Lupinus Albus | 0.57±0.057 | Oats | 1.6±0.163 |
| Cr(mg/kg) | 2.6±0.26 | Wheat | 1.3±0.13 | White(barley,Teff) | 1.8±0.171 |

### Concentration of essential elements

Based on **Table 5**, the mean concentrations (mean ± SD, n = 3) of Fe, Zn, Se, Mn, Cu and Mo varied significantly among the analyzed Ethiopian herbal food supplements. Iron (Fe) ranged from (68.6 ± 6.9) mg/kg (chickpea) to (506.5 ± 50.7) mg/kg (wheat). Zinc (Zn) ranged from (1.3 ± 0.13) mg/kg in chickpea to (8.1 ± 0.81) mg/kg in *Lupinus albus*; selenium (Se) ranged from (0.026 ± 0.003) µg/kg in chickpea to (0.095 ± 0.01) µg/kg in *Lupinus albus*; manganese (Mn) ranged from (4.0 ± 0.4) mg/kg in chickpea to (183.9 ± 18.4) mg/kg in *Lupinus albus*; and copper (Cu) ranged from (1.6 ± 0.16) mg/kg in white barley to (4.6 ± 0.5) mg/kg in wheat. Manganese (Mn) ranged from (4.0 ± 0.4) mg/kg (chickpea) to (183.9 ± 18.4) mg/kg (*Lupinus albus*), whereas selenium (Se) ranged from (0.026 ± 0.003) µg/kg (chickpea) to (0.095 ± 0.01) µg/kg (*Lupinus albus*). Molybdenum (Mo) varied from (0.064 ± 0.006) µg/kg (black barley) to (4.4 ± 0.044) µg/kg (*Lupinus albus*), whereas copper (Cu) ranged from (1.6 ± 0.16) mg/kg (white barley) to (4.6 ± 0.5) mg/kg (wheat).

**Table 5:** the mean amounts (mean ± SD) of the metals Fe (mg/kg), Zn (mg/kg), Se (µg/kg), Mn (mg/kg), Cu (mg/kg), and Mo (µg/kg) in Ethiopian herbal food supplements (n = 3)

| Food samples | Trace elements |  |  |  |  |  |
| --- | --- | --- | --- | --- | --- | --- |
| | Fe (mg/kg) | Zn (mg/kg) | Se (μg/kg) | Mn (mg/kg) | Cu (mg/kg) | Mo (μg/kg) |
| Wheat | 506.5±50.7 | 11.2±1.1 | 0.033±0.003 | 23.5±2.4 | 4.6±0.5 | 0.471±0.047 |
| Chickpea | 68.6±6.9 | 1.3±0.13 | 0.026±0.003 | 4.0±0.4 | 0.597±0.06 | 0.132±0.013 |
| Oats | 404.4±40.4 | 4.6±0.46 | 0.088±0.009 | 14.4±1.4 | 1.9±0.2 | 0.206±0.021 |
| TeffWhite | 144.0±14.4 | 3.6±0.36 | 0.044±0.004 | 18.2±1.8 | 1.9±0.2 | 0.130±0.013 |
| TeffRed | 291.8±29.2 | 7.1±0.71 | 0.081±0.008 | 43.5±4.4 | 3.1±0.31 | 0.361±0.036 |
| BlakBarley | 161.3±16.1 | 3.4±0.34 | 0.056±0.006 | 37.6±3.8 | 2.2±0.22 | 0.064±0.006 |
| WhiteBarley | 92.0±9.2 | 3.6±0.36 | 0.034±0.003 | 12.1±1.2 | 1.6±0.16 | 0.152±0.015 |
| Lupinus albus | 312.2±31.2 | 8.1±0.81 | 0.095±0.01 | 183.9±18.4 | 4.2±0.42 | 4.4±0.044 |
| Pumpkin | 319.2±31.9 | 6.0±0.6 | 0.058±0.006 | 150.9±15.1 | 3.5±0.35 | 0.058±0.006 |

The highest, minimum, and overall average concentrations of Fe, Zn, Se, Mn, Cu, and Mo throughout the examined herbal food supplements are summarized in (**Table 6)**, which highlights the variation among food types and serves as a foundation for nutritional and safety assessment[14].

**Table 6:** East Gojjam herbal food supplements’ maximum, minimum, and average concentrations of Fe (mg/kg), Zn (mg/kg), Se (µg/kg), Mn (mg/kg), Cu (mg/kg), and Mo (µg/kg) (mean± SD) (n = 3)

| Radionuclides | Maximum | Sample Type | Minimum | Sample type | Average |
| --- | --- | --- | --- | --- | --- |
| Fe(mg/kg) | 506.5±50.7 | Wheat | 68.6±6.9 | Chickpea | 280.7±25.6 |
| Zn(mg/kg) | 11.2±1.1 | Wheat | 1.3±0.13 | Chickpea | 6.0±0.541 |
| Se (μg/kg) | 0.095±0.01 | *lupinus albus* | 0.026±0.003 | Chickpea | 0.061±0.006 |
| Mn(mg/kg) | 183.6±18.4 | *lupinus albus* | 4±0.4 | Chickpea | 398.3±23.8 |
| Cu(mg/kg) | 4.6±0.5 | Wheat | 1.6±0.16 | BarleyWhite | 2.8±0.269 |
| Mo (μg/kg) | 4.4±0.44 | *lupinus albus* | 0.064±0.006 | BarleyBlack | 1.0±0.022 |

#### Comparative Analysis

Nine herbal dietary supplements have a total of six elemental impurities. The concentrations of the important metals (Fe, Cu, Mn, Mo, Se, and Zn) in Ethiopian herbal dietary supplements (n = 3) are shown in (**Table 7)** together with published values from other nations[40]. Fe, Cu, Mn, and Zn are reported in mg/kg, while Mo and Se are provided in µg/kg when available. The concentrations were expressed on a dry weight basis and are shown in the relevant units used throughout investigations[14]. This makes it possible to meaningfully compare the food samples from East Gojjam with published data from various geographical locations [41].

**Table 7.**
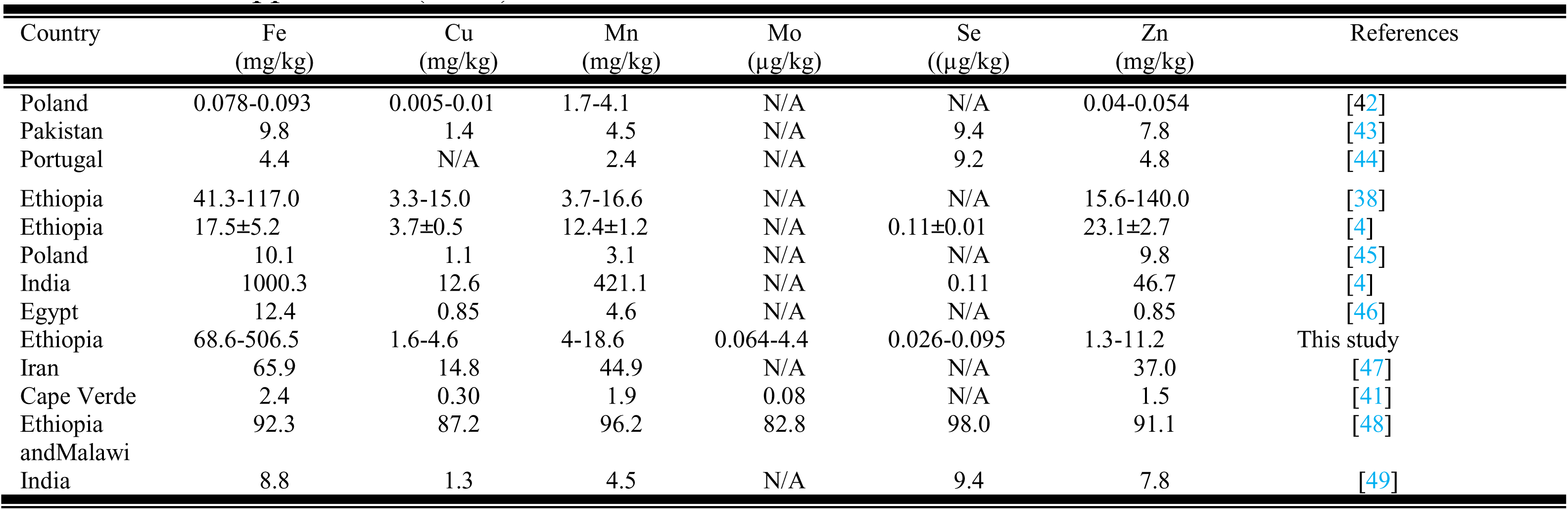
Comparison of activity concentration of Fe, Cu, Mn, Mo, Se, Zn of Ethiopian herbal food supplements (n = 3) with their values for various nations worldwide.

Comparing concentrations across nations offers valuable information for nutritional profiling, indicating the potential nutritional contribution of East Gojjam herbal food supplements with respect to important dietary micronutrients like iron, zinc, copper, manganese, selenium, and molybdenum [41]. Significant variation in the measured essential element levels can be attributed to differences in plant species and variety, soil geochemistry, environmental conditions, agricultural practices, and analytical methodology (including digestion conditions and ICP-MS calibration)[14]. In order to determine the potential nutritional contribution of East Gojjam herbal food supplements with regard to important dietary micronutrients like iron, zinc, copper, manganese, selenium, and molybdenum, it is helpful to compare concentrations across nations [41].

Overall, **Table 7’s** inter-country comparison offers context for assessing the nutritional quality of the investigated herbal food supplements and supports the evaluation of essential micronutrient availability[38].

### Overview of the studied herbal food supplements

The studied herbal foods supplements: teff (*Eragrostis teff*), lupin (*Lupinus albus*), oats (*Avena sativa*), barley (*Hordeum vulgare*), pumpkin/*cucurbita* (*Cucurbita spp*.), wheat (*Triticum aestivum*), and chickpea (*Cicer arietinum*)—are commonly consumed cereal/non-cereal seeds in East Gojjam. Teff, oats, barely and wheat have been produced for nourshiment in various traditional preparations, including porridge, soups, stews, flatbread (injera), snacks, local bear (Tella) and flour-based foods in East Gojjam [38,40]. Their nutritional contributions depend on their macro- and micronutrient composition, including essential trace elements[38].

Because these foods may provide dietary essential elements while also containing trace contaminants depending on soil and processing conditions, their elemental composition was determined using ICP-MS in this study [40].

### General Discussion on Nutritional values and elemental profiling

The present ICP-MS analysis quantified essential elements (Fe, Zn, Se, Mn, Cu and Mo) in East Gojjam herbal food supplements to support both nutritional evaluation and potential safety considerations [30].Essential trace elements contribute to key physiological functions, and their intake depends on both the concentration in foods and dietary consumption patterns[40].

Overall, iron (Fe) showed the greatest concentration in wheat (506.5±50.7) mg/kg, while the lowest Fe level was observed in chickpea (68.6±6.9) mg/kg. Zinc (Zn) varied markedly, ranging from (1.3±0.13) mg/kg in chickpea to (8.1±0.81) mg/kg in *Lupinus albus*. Selenium (Se) was detected at low levels (µg/kg range), with higher values in *Lupinus albus* (0.095±0.01) µg/kg compared with chickpea (0.026±0.003) µg/kg. Manganese (Mn), copper (Cu) and molybdenum (Mo) also displayed clear variability among the samples, indicating that different cereal and non-cereal seeds may contribute differently to micronutrient intake[50].

From a nutritional perspective, foods such as wheat and *Lupinus albus* may provide relatively higher contributions of Fe and Zn, respectively, while pumpkin and oats showed intermediate profiles depending on the specific element[33,35]. Such elemental variation supports the importance of ICP-MS-based elemental profiling when recommending dietary intake of herbal foods, particularly where these foods are used as staple components [38].

## Conclusions

This study used ICP-MS to quantify both nutritional essential elements (Cu, Fe, Zn, Mo, Se, Mn) and trace toxicants (notably Pb, Al, Ni, and Cr) in herbal food supplement matrices following microwave acid digestion with 8 mL (HNO₃,70%) + 2 mL (H₂O₂,30%). The analytical performance of the method was supported by robust validation: calibration linearity was high with R^2^ ranging from (0.9993-0.9999), standards were re-run using the same sample calibration, CRM recoveries (88.9–110%) demonstrated good trueness, and within-run RSDs were <5% for most analytes, indicating good precision and reliability of the concentration data used for subsequent health-risk computations.

Analytical sensitivity was adequate for trace-level profiling. The target analytes’ LOD ranges were (0.001–0.01) mg/kg for Cu, Fe, Zn, Mn, Ni, and Cr and (0.002–0.05) mg/kg for Mo, Se, Pb, and Al. Accordingly, the LOQs varied from 0.01 ppm (Cr) to 0.265 ppm (Fe), allowing for accurate measurement of both toxicant and nutritious constituents in the supplement matrices.

HQ/HI was used to evaluate non-carcinogenic risk; for all samples, HQ < 1 meant that the consumption of each non-carcinogenic toxicant in the products did not surpass the threshold for possible non-carcinogenic consequences. Even though single-element HQ values remained below 1, the hazard index (HI) for *Lupinus albus* reached 1.4, suggesting that the cumulative exposure to the evaluated elements in this product may represent a possible non-carcinogenic issue.

Carcinogenic risk was evaluated using ILCR. Overall, ILCR values were < 10^−4^ for all samples, indicating a generally low carcinogenic risk across the product set. In contrast, for *Lupinus albus* the ILCR increased to > 10^−3^ and was specially attributed to nickel (Ni). This element-driven elevation is consistent with the reported element-specific ILCR ranges: Pb: (9.7×10^−9^ to 1.2×10^−7^), Cr: (1.9×10^−4^ to 4.1×10^−4^), and Ni: (1.4×10^−4^ to 1.1×10^−3^).

Nutritional profiling confirmed that the supplements contained measurable essential nutrients (Cu, Fe, Zn, Mo, Se, Mn), which supports their potential nutritional contribution. Nevertheless, nutritional benefit should be interpreted alongside co-occurrence of toxic elements.

The ICP-MS method was analytically valid and sensitive (supported by LOD/LOQ, validation parameters,CRM recoveries, and RSD precision). From a toxicological perspective, non-carcinogenic risk was generally low (HQ < 1), but combined exposure in *Lupinus albus* may be elevated (HI = 1.4). Carcinogenic risk was also generally low (ILCR < 10^−4^), except that *Lupinus albus* exhibited higher carcinogenic risk (ILCR > 10^−3^) largely due to Ni. Therefore, *Lupinus albus* should be prioritized for tighter elemental quality control, particularly for nickel.

## Future Developments

Similar to other agro-food systems, cereal and non-cereal researches necessitate a systemic approach involving collaboration among the agricultural research community and national and international organizations.

## The authors’ contribution

AYH organized, executed, and completed the statistical and laboratory analyses, sampling, and sample preparation. MM was in charge of the entire project, edited the material, made important adjustments, and assisted with setting up the experiment.The final draft was examined and approved by each author.

## Data availability

Upon request, the relevant author will supply the data used to bolster the study’s results.

## Acknowledgment

The kind hospitality, counsel, direction, and participation of the people of East Gojjam, Ethiopia, were essential to the success of this study.

